# Unique features of magic spot metabolism in *Clostridioides difficile*

**DOI:** 10.1101/2021.08.02.454818

**Authors:** Asia Poudel, Astha Pokhrel, Adenrele Oludiran, Estevan J. Coronado, Kwincy Alleyne, Marrett M. Gilfus, Raj K. Gurung, Surya B. Adhikari, Erin B. Purcell

## Abstract

The ‘magic spot’ alarmones (pp)pGpp, previously implicated in *Clostridioides difficile* antibiotic survival, are synthesized by CdRSH and CdRelQ. These enzymes are transcriptionally activated by diverse environmental stresses, but both exclusively synthesize pGpp rather than ppGpp as has previously been reported. While direct synthesis of pGpp from a GMP substrate and (p)ppGpp hydrolysis into pGpp by NUDIX hydrolases have previously been reported, there is no precedent for a bacterium synthesizing pGpp exclusively. Hydrolysis of the 5’ phosphate or pyrophosphate from GDP or GTP substrates is necessary for activity by the clostridial enzymes, neither of which can utilize GMP as a substrate. Both enzymes are remarkably insensitive to the size of their metal ion cofactor, tolerating a broad array of metals that do not allow activity in (pp)pGpp synthetases from other organisms. It is clear that while *C. difficile* utilizes magic spot signaling, its mechanisms of alarmone synthesis are not directly homologous to those in more completely characterized organisms.

## Introduction

*Clostridioides* (formerly *Clostridium*) *difficile* is a Gram positive, spore forming anaerobic gastrointestinal pathogen that causes *Clostridioides difficile* infection (CDI), which is multidrug resistant and has a high rate of recurrence ^1^ *C. difficile* is transmitted to susceptible hosts via faeco-oral transmission. ^2^ Once spores are ingested, they survive the acidic pH of the stomach and travel through the intestine where they interact with primary bile salts and amino acids that facilitate their germination into vegetative cells which secrete the toxins that cause symptomatic disease. ^3^ The most significant risk factor for CDI development is the exposure of the normal gut microflora to broad spectrum antibiotics. ^1, 4^

*C. difficile* must navigate a dynamic and hostile environment to establish symptomatic infection. The proliferation of vegetative cells in the gut environment is impacted by several factors including the host immune response, oxygen levels, nutrient limitation, secondary bile acids, short chain fatty acids, competing gut microflora and varying pH throughout the gastrointestinal tract.^5–11^ The immune response to gastrointestinal infection can include fever of 1-4°C, antimicrobial host defense peptides, sequestration of metals to limit their availability, and oxidative stress from reactive oxygen species (ROS) and reactive nitrogen species (RNS).^12–17^

The stringent response (SR) is a stress survival mechanism that enables bacterial cells to quickly adapt to extracellular stress by temporarily halting growth and cell division and inducing transcription of stress survival genes. ^18–19^ The SR contributes to regulation of bacterial growth, virulence, antimicrobial peptide survival, oxidative stress resistance, antibiotic tolerance and persistence in diverse bacterial pathogens. ^20^ This process is canonically driven by the accumulation of two intracellular alarmone nucleotides, guanosine pentaphosphate (pppGpp) and guanosine tetraphosphate (ppGpp) collectively known as (p)ppGpp and sometimes referred to as ‘magic spot’. ^21^ These signaling molecules are metabolized by multidomain bifunctional synthetase/hydrolase enzymes in the long RelA-SpoT (RSH) family and by monofunctional small alarmone synthetase (SAS) and small alarmone hydrolase (SAH) domains.^22^ Extracellular stresses including amino acid starvation, alkaline shock, cell wall stress from cell wall active antibiotics, heat stress, oxidative stress and acid stress can trigger the SR modulating the transcription of either RSH or SAS genes.^18, 23, 24, 25–26^ Post-transcriptional regulation of RSH enzymes typically involves protein-protein interactions or binding of uncharged tRNAs, while post-transcriptional regulation of SAS enzyme seems to depend on allosteric binding of (p)pGpp. ^27–30^ The SAS enzymes also appear to play a stress-independent role in maintaining basal (p)ppGpp levels and regulating guanosine homeostasis in *Bacillus subtilis* and *E. faecalis*.^31–33^

Guanosine substrate utilization varies among characterized alarmone synthetases, hinting at different biological roles. We have previously reported that CdRSH binds both GDP and GTP *in vitro* but exclusively utilizes GDP as a phosphoacceptor in order to synthesize ppGpp.^34^ This is unusual among RSH family enzymes, which utilize both GDP and GTP with varying affinity to synthesize ppGpp and pppGpp and typically have higher affinity for GTP in Gram positive organisms ^22, 35–39^ In addition, RSH homologs from *B. subtilis and Methylobacterium exotorquens* have recently been reported to synthesize (p)ppApp when incubated with adenosine phosphoacceptors. ^40–41^ The RelP and RelQ enzymes of *B. subtilis*, *E. faecalis*, and *S. aureus* all exhibit greater affinity for GDP than GTP. ^31, 42–43^ The RelP and RelQ enzymes from *S. mutans*, while active *in vivo,* have not been purified for *in vitro* study. ^44^ RelZ from *M. smegmatis*, which contains an RNaseH domain in addition to the SAS, utilizes GMP preferentially. ^45^ RelS from *C. glutamicum,* part of a subfamily found only in *Acinetobacter*, has a higher affinity for GTP. ^46^ *C. glutamicum* also contains a RelP homolog, RelP*, which lacks conserved amino acids at its C-terminus and has no (pp)pGpp synthetic capacity *in vivo* ^46^.

Firmicutes species typically encode a single RSH and one or two SAS.^22^ The highly conserved synthetase domains of both families catalyze the transfer of a pyrophosphate moiety from ATP to 3^’^-OH of GDP and/or GTP to produce ppGpp or pppGpp, respectively.^21^ SAS enzymes from several Gram positive species were recently discovered to synthesize a third alarmone, pGpp, utilizing GMP as a substrate.^31, 46–47^ In addition, the recently discovered NUDIX hydrolases remove 5’ phosphates or pyrophophates from (p)ppGpp to produce pGpp.^31, 45, 47–49^ The physiological role of pGpp as a third alarmone is still under investigation. It has been speculated to allow a prolonged SR after the depletion of cytoplasmic GTP and GDP.^31, 46^ We will henceforth refer to the three alarmones collectively as (pp)pGpp where appropriate.

Here, we report that expression of clostridial (pp)pGpp synthetase genes is induced by antibiotic and acid stress. We further report that the RelQ enzyme utilizes both GDP and GTP as substrates, but that the only form of alarmone produced by either CdRSH or CdRelQ enzyme is pGpp. Activity of both enzymes depends upon 5’ beta phosphate bond hydrolysis of the GXP substrate. We find that CdRelQ, like CdRSH, tolerates a remarkable diversity of divalent metal ion cofactors. The seemingly exclusive production of pGpp is unique to *C. difficile* among previously studied bacteria.

## RESULTS

### C. difficile encodes one conserved SAS and one divergent SAS

We have previously reported that *C. difficile* encodes an RSH family enzyme and the SAS RelQ, and confirmed the enzymatic activity of RSH. ^34^ In addition to RelQ, *C. difficile* encodes another putative SAS enzyme, CDR20291_1607, which we have named RelC. RelC contains a putative SAS domain and a C-terminal 250 residue region that contains no conserved domain, and its only homologs are in *Firmicutes* species. ^22^ CdRelQ exhibits high sequence conservation in the ATP and GDP binding motifs identified in crystal structures of *B. subtilis* RelQ and *S. aureus* RelP (Figure 1). ^37, 50^ CdRelC has a highly conserved ATP-binding motif, but the putative GDP-binding motif lacks several conserved residues including a tyrosine (Y116 in BsRelQ, Y151 in SaRelP, Q137 in RelC) that stacks with the guanosine base in the GDP substrate and is invariant in SAS and RSH synthetase domains. ^37, 50^ In addition, RelC contains a ten residue insert within the GDP recognition motif that has no homolog in any characterized SAS (Figure 1). Initial attempts to express RelC in *E. coli* for purification were unsuccessful, but further experimentation will be necessary to determine whether RelC binds any form of guanosine substrate or serves as a pyrophosphotransferase. (Figure 1).

**Figure 1.**
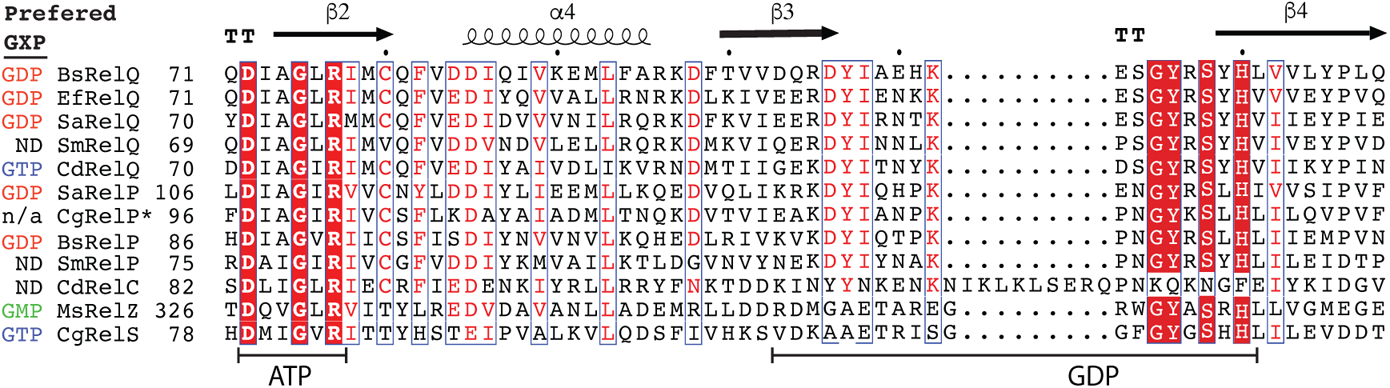
Substrate binding sites of SAS enzymes. The sequences of characterized SAS enzymes are aligned with the crystal structure of *Bacillus subtilis* RelQ (PDB 5DED). β-strands are shown as arrows and α-helices are shown as corkscrews. Conserved residues are shown in red and highly conserved residues are shown in white on a red background. The ATP and GDP binding motifs identified in BsRelQ and SaRelP are indicated on the bottom. The preferred guanosine nucleotide substrate of enzymes that have been kinetically characterized are shown on the left. ND indicates that the substrate preference of an enzyme has not been determined; n/a indicates that the enzyme is not active as a (pp)pGpp synthetase. Alignment generated with ESPript 3 (https://espript.ibcp.fr/ESPript/ESPript/).

### C. difficile induces rsh and relQ transcription in response to clinically relevant antibiotic stress

In our earlier work, we utilized an anaerobic fluorescent transcriptional reporter incorporating the oxygen-independent fluorescent flavoprotein phiLOV2.1 to monitor transcription of the clostridial *rsh* and *relQ* genes.^34^ We observed the increased transduction of these genes upon exposure to sublethal concentrations of clindamycin and metronidazole in a strain-specific manner.^34^ Here, we assessed the transcriptional response of *C. difficile* to vancomycin and fidaxomicin, the currently recommended antibiotic treatments for severe or recurring CDI.^51–53^ Exponentially growing cells were exposed to drug concentrations 0.5x of those sufficient to inhibit growth (Figure 2A). In the epidemic strain *C. difficile* R20291 promoter activity in the tetracycline-inducible control strain was unaffected by either antibiotic. Vancomycin had no effect on transcription from the *rsh* or *relQ* promoters. Fidaxomicin stimulated a 96% increase in P*_rsh_* activity and a 3.1-fold increase in P*relQ* activity (Figure 2B). In the laboratory strain *C. difficile* 630Δ*erm*, the P*_rsh_* reporter had no response to either drug, while activity from the P*_relQ_* reporter increased 3.6-fold upon exposure to vancomycin (Figure 2C).

**Figure 2.**
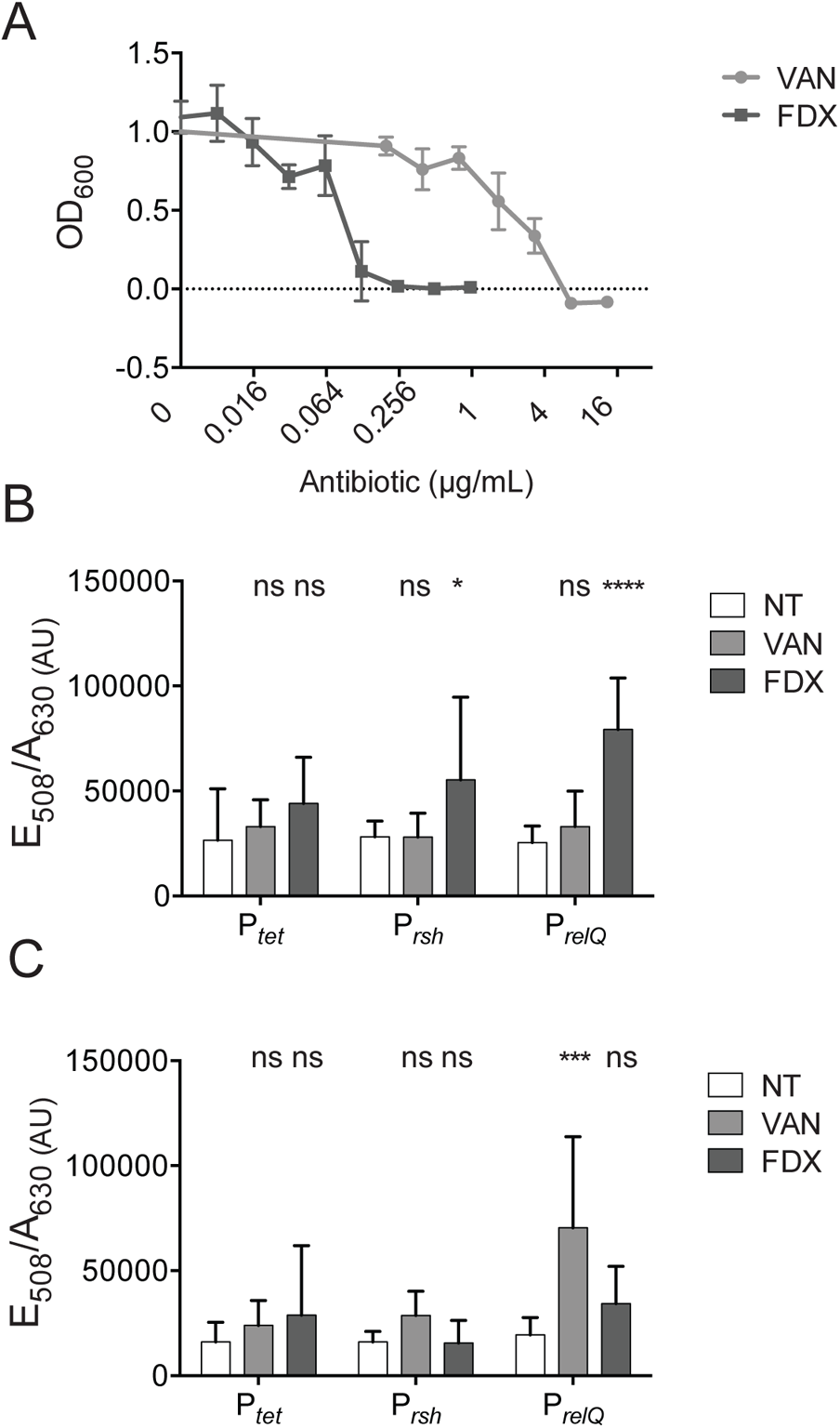
Transcriptional response of (pp)pGpp synthetase genes to antibiotic stress. (A) Overnight growth of R20291 in the indicated concentrations of vancomycin (VAN) and fidaxomicin (FIX). Shown are the means and standard deviations of nine biologically independent samples. (B and C) PhiLOV fluorescence normalized to cell density after two hours of exposure to 1.7 µg mL^−1^ vancomycin or 0.067 µg mL^−1^ fidaxomicin. Shown are the means and standard deviations of at least six biologically independent samples in the (B) R20291 and (C) 630Δ*erm* backgrounds. Conditions treated with antibiotics are compared to untreated by two-way ANOVA. **** *p* < 0.0001, * *p* < 0.05, ns not significant.

### rsh and relQ transcription during oxidative, heat, and pH stress

To assess the potential role of (pp)pGpp signaling and the stringent response in *C. difficile* survival of environmental stresses within the host, we monitored *rsh* and *relQ* promoter activity in *C. difficile* R20291 under conditions mimicking potential stressors *in vivo*. Activity from both promoters was indistinguishable after incubation at either 37°C or 41°C, suggesting that even high fever within a mammalian host would not activate transcription of (pp)pGpp synthetase genes (Figure 3A). Oxidative stress was mimicked through exposure to copper, a transition metal previously shown to inhibit *C. difficile* growth, and to diamide, an oxidant which mimics oxidative stress in anaerobes by instigating disulfide bonds. ^54–55^ Three hours of exposure to copper did not stimulate increased activity from the *rsh* or *relQ* promoters; indeed, *rsh* promoter activity was somewhat decreased after copper treatment (Figure 3C). Treatment with diamide stimulated a 3.8 fold increase in *rsh* promoter activity but had no effect on the *relQ* promoter (Figure 3D). While the baseline fluorescence of the P*_tet_* reporter strain increased in alkaline conditions, transfer to media buffered at neutral or alkaline pH had no effect on the activity of either clostridial promoter relative to that of the of the P*_tet_* reporter (Figure 3E). However, transfer from unbuffered BHIS (initial pH measured at 7.4) to BHIS buffered at pH 6 did stimulate a 3.2 fold increase in P*_rsh_* activity (Figure 3E). Notably, when we measured the pH of spent BHIS after 24 hours of *C. difficile* growth, we found that R20291 acidifies its growth medium to pH 6. Medium acidification by 630Δ*erm* is less pronounced (Table 1). We have previously shown that *rsh* transcription in *C. difficile* is induced by stationary phase onset.^34^ It is possible that medium acidification, in addition to nutrient depletion, triggers this response.

**Figure 3.**
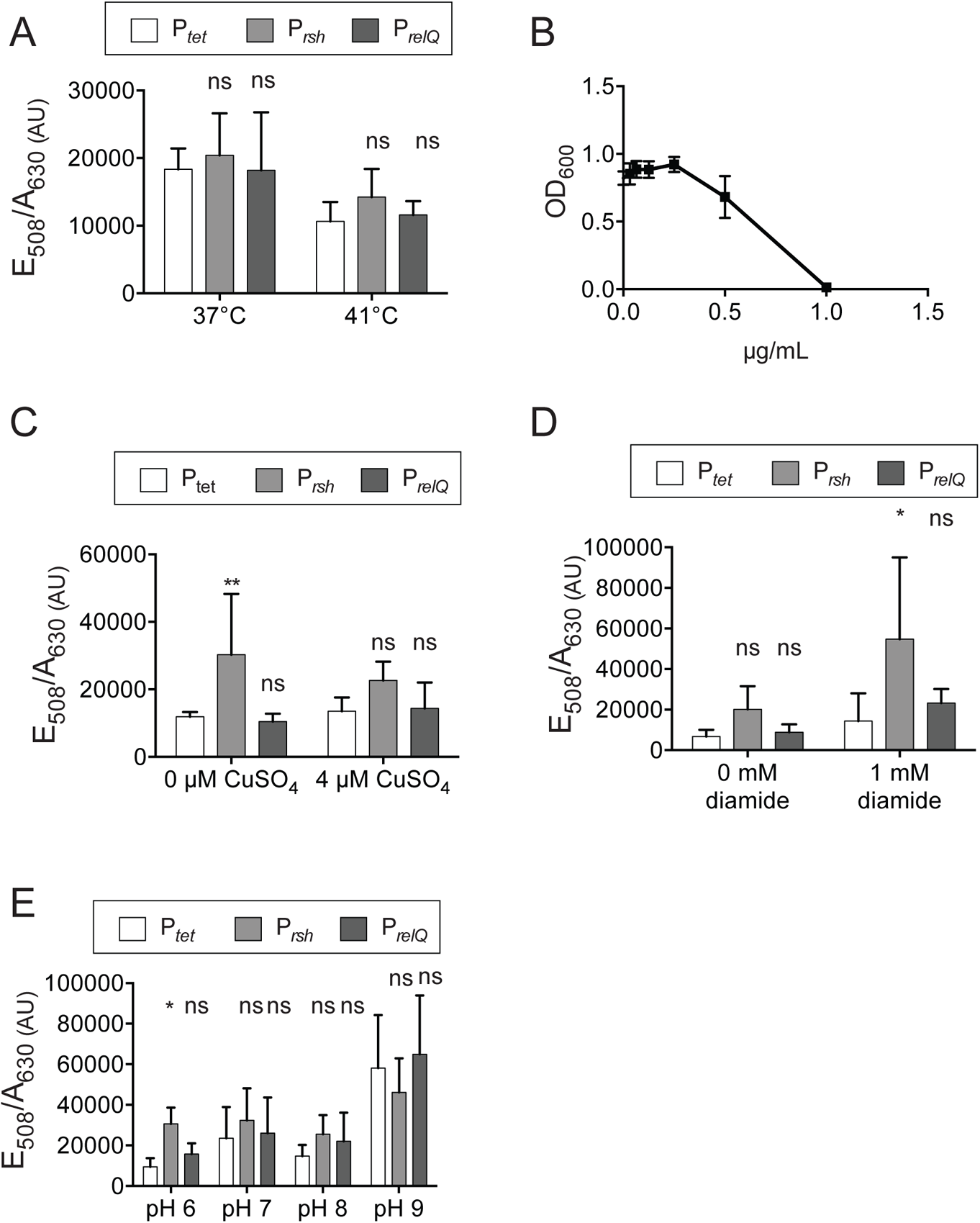
Transcriptional response of (pp)pGpp synthetase genes to environmental stress in R20291. (A) PhiLOV fluorescence normalized to cell density after 30 minutes of incubation at the indicated temperature. Shown are the means and standard deviations of six biologically independent samples. (B) Cell density after overnight growth in TY medium containing the indicated concentration of diamide. Shown are the means and standard deviations of three biologically independent samples. (C) PhiLOV fluorescence normalized to cell density after two hours of exposure to the indicated concentration of copper sulfate. Shown are the means and standard deviations of six biologically independent samples. (D) PhiLOV fluorescence normalized to cell density after three hours of exposure to the indicated concentration of diamide. Shown are the means and standard deviations of three biologically independent samples. PhiLOV fluorescence normalized to cell density three hours after transfer from unbuffered BHIS to BHIS buffered at the indicated pH with potassium phosphate buffer. (A, C-E) Activity from the P*rsh* and P*relQ* promoters was compared to that from the P*tet* promoter in the same condition by two-way ANOVA. * *p* < 0.05, ** *p* < 0.01, ns not significant.

**Table 1.** pH of fresh and spent BHIS medium. pH of growth medium was measured before and 24 hours after inoculation with single colonies.

| *C. difficile* strain | pH |  |
| --- | --- | --- |
|  | Fresh BHIS | 24 h spent BHIS |
| R20291 | 7.44 | 6.06 |
| 630Δ*erm* | 7.44 | 6.94 |

### CdRelQ is an active (pp)pGpp synthetase

We have previously reported the *in vitro* synthetase activity of RSH from *C. difficile*. Here, we confirm that the putative synthetase RelQ from this organism is also active *in vitro*, capable of transferring a pyrophosphate moiety from γ-^32^P-ATP to a guanosine nucleotide phosphoacceptor (Figure S1A). Unlike CdRSH, CdRelQ activity is concentration dependent and exhibits little to no pyrophosphotransfer activity at concentrations below 0.4 µM (Figure S1A,B). This may explain why, unlike CdRSH, CdRelQ does not arrest cell division when overexpressed in *E. coli*, suggesting a lack of *in vivo* (pp)pGpp synthesis activity (Figure S1C). SAS from *B. subtilis*, *S. aureus*, and *E. faecalis* function as tetramers, raising the possibility that CdRelQ is not expressed in *E. coli* at levels sufficient to allow oligomerization and activity. ^37, 50, 56^

### CdRelQ utilizes GDP and GTP but synthesizes pGpp exclusively

In order to assess the substrate specificity of CdRelQ, we performed synthesis assays utilizing the monophosphate, diphosphate, and triphosphate forms of both adenosine and guanosine nucleotides as phosphoacceptors. We found that CdRelQ is incapable of transferring pyrophosphate from ATP to any adenosine nucelotides or to guanosine monophosphate (Figure 4A). CdRelQ utilizes both GDP and GTP as phosphoacceptors, but synthesizes the same size product rather than the expected ppGpp and pppGpp, which should migrate differently on a chromatogram due to the additional mass and charge of the pentaphosphate alarmone (Figure 4A). We performed synthesis assays using the same range of adenosine and guanosine phosphoacceptors with the BsRelQ enzyme from *Bacillus subtilis*, which is known to utilize GMP, GDP, and GTP *in vitro* in order to synthesize pGpp, ppGpp, and pppGpp, respectively.^33, 40^ As predicted, BsRelQ utilized the three guanosine nucleotide substrates to synthesize three differently sized products. Unexpectedly, the single product synthesized by CdRelQ had the same apparent size and charge as pGpp synthesized by BsRelQ from ATP and GMP, rather than ppGpp (Figure 4A). Upon further exploration of guanosine metabolism utilization by these enzymes, CdRSH utilized GDP to synthesize an apparent pGpp product. In this assay, CdRSH exhibited some trace synthesis of an apparent pGpp product using GTP as a substrate, although the pGpp spot was very faint and could have resulted from GDP contamination in the GTP sample (Figure 4B). BsRelQ again utilized mono-, di-, and triphosphate guanosine substrates to synthesize tri-, tetra-, and pentaphosphate products, respectively. CdRelQ utilized both GDP and GTP as substrates but only synthesized the apparent triphosphate pGpp product. Neither clostridial enzyme utilized GMP as a substrate and neither synthesized any product other than the apparent triphosphate magic spot pGpp. To rule out the possibility that the clostridial enzymes synthesize (p)ppGpp and then convert it to pGpp through an intrinsic hydrolysis capability, we incubated CdRelQ with ppGpp and pppGpp produced by BsRelQ. After an hour of incubation, the exogenously produced magic spots had not been hydrolyzed and no pGpp spot had emerged, suggesting that CdRelQ synthesizes pGpp directly rather than converting (p)ppGpp intermediates (Figure 4C).

**Figure 4.**
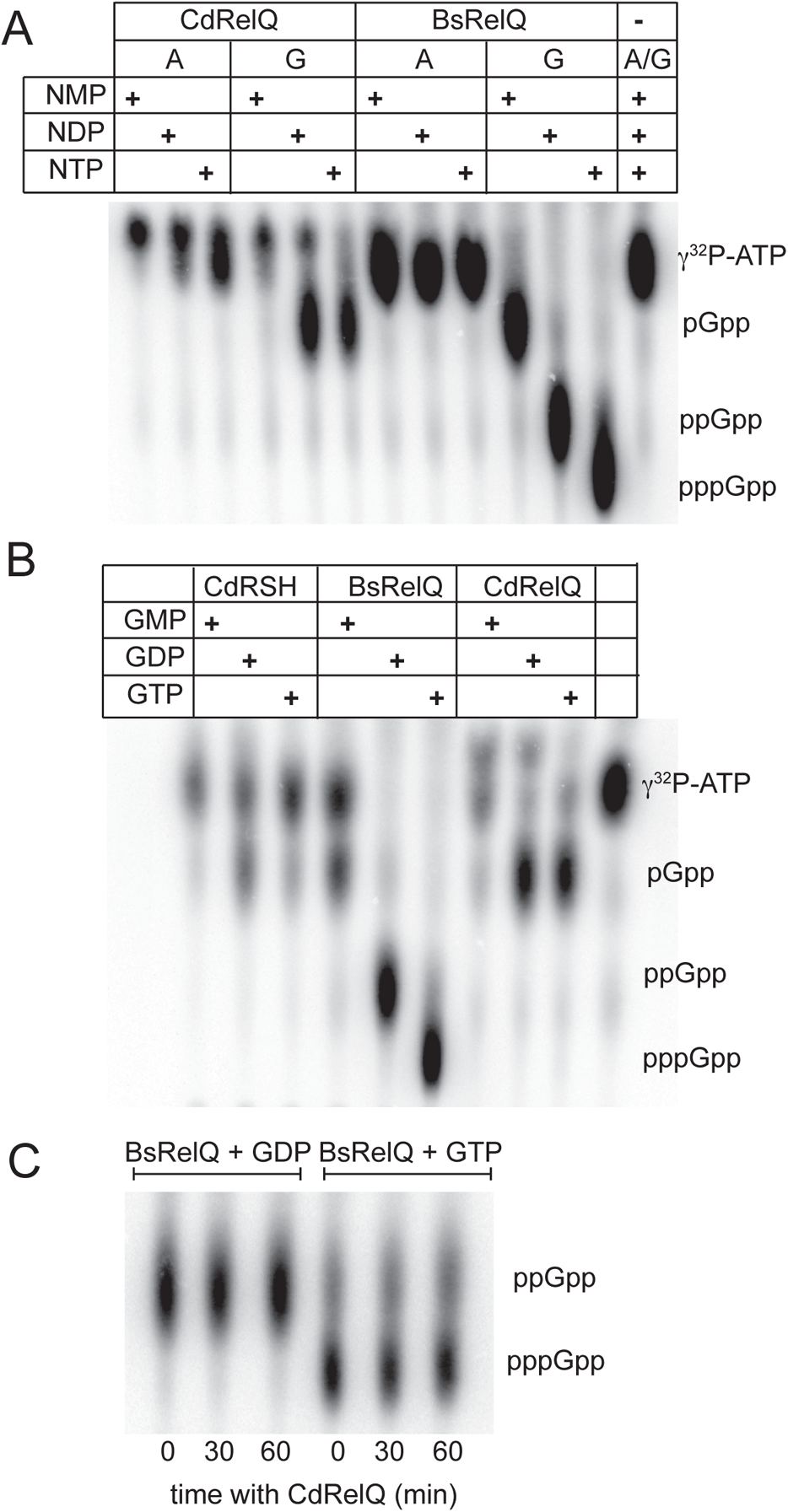
CdRelQ utilizes GDP and GTP but exclusively synthesizes pGpp. (A) Pyrophosphotransfer after 60 minutes from γ-^32^P-ATP to 500 µM of the indicated purine nucleotide phosphoacceptors by 2.0 µM CdRelQ or 3.0 µM BsRelQ. (B) Pyrophosphotransfer after 60 minutes from γ-^32^P-ATP to 500 µM of the indicated guanosine nucleotide phosphoacceptors by 3.0 µM CdRSH, 3.0 µM BsRelQ, or 2.0 µM CdRelQ. (C) After BsRelQ was incubated with GDP or GTP for 60 minutes, CdRelQ was added to the reaction for the indicated time and reaction products were analyzed by TLC..

Under the conditions studied, CdRelQ appeared to consume all of the γ-^32^P-ATP provided in the presence of GTP but leave more residual ATP in the presence of GDP, suggesting a higher affinity for the triphosphate substrate (Figure 4A). Kinetic analysis of enzyme activity at multiple concentrations of guanosine substrate yielded calculated Michaelis constants of 45.4 µM for GTP and 53.6 µM for GDP, consistent with the visual observations (Table 2, Figure S1). In order to further quantitate the affinity of CdRelQ for its guanosine substrate we performed isothermal titration calorimetry. We found that the affinity constant of CdRelQ for GDP is roughly half that for GTP (Table 2). While the two results did not produce a consensus for the absolute binding affinity of RelQ for its guanosine substrates, they were consistent in indicating the enzyme has higher affinity for the triphosphate substrate under the experimental conditions. As CdRSH has a higher affinity for GDP than GTP *in vitro*, these results suggest that the two synthetases may be active in discreet cellular conditions when either GTP or GDP are more prevalent.

**Table 2.** Substrate affinity of CdRelQ at 37°C. The binding of CdRelQ at 0.006 mM to GDP/GTP at 0.060 mM was measured by ITC. The data were fitted to a single-binding site model. Shown are the values for K, the equilibrium binding constant; ΔH, the enthalpy change associated with protein binding to the ligand; and ΔS, the entropy change associated with binding. Each value is the average of two repeat experiments and the standard deviation ± are shown

| Michaelis-Menton analysis |  |  |  |
| --- | --- | --- | --- |
| Substrate | $K_M$ ($M^{-1}$) | | |
| GDP | 45.4 |  |  |
| GTP | 53.6 |  |  |
| Isothermal titration calorimetry |  |  |  |
| Substrate | $K$ ($M^{-1}$) | $\Delta H$ (cal mol$^{-1}$) | $\Delta S$ (cal mol$^{-1}$ °C$^{-1}$) |
| GDP | $2.04E3 \pm 2.25E3$ | $-8.18E7 \pm 8.21E7$ | $-2.64E5$ |
| GTP | $4.87E4 \pm 3.64E3$ | $-1.639E6 \pm 7.412E4$ | $-5.25E3$ |

### Clostridial synthetases require phosphorolysis of the guanosine substrate

To confirm that the guanosine substrate is hydrolyzed during clostridial magic spot synthesis, we employed 5’-β-thio-diphosphate (GDPβS), which has a non-hydrolyzable thiol bond between the α and β phosphates, as a phosphoacceptor. Both CdRelQ and CdRSH synthesized pGpp when GDP was supplied as a substrate, but neither clostridial enzyme generated any form of magic spot when GDPβS was the substrate (Figure 5). RelQ did appear to deplete the radioactive ATP substrate in the presence of GDPβS but could not complete phosphotransfer, while RSH displayed no activity at all in the absence of a chemically labile β-phosphate bond on the guanosine substrate (Figure 5). BsRelQ was able to utilize GDPβS as a phosphoacceptor. The product formed by BsRelQ using GDPβS is larger in size than ppGpp formed from GDP since the non-hydrolysable analogue contains three lithium atoms that make the product heavier than ppGpp and closer in apparent size to pppGpp formed from GTP (Figure 5).

**Figure 5.**
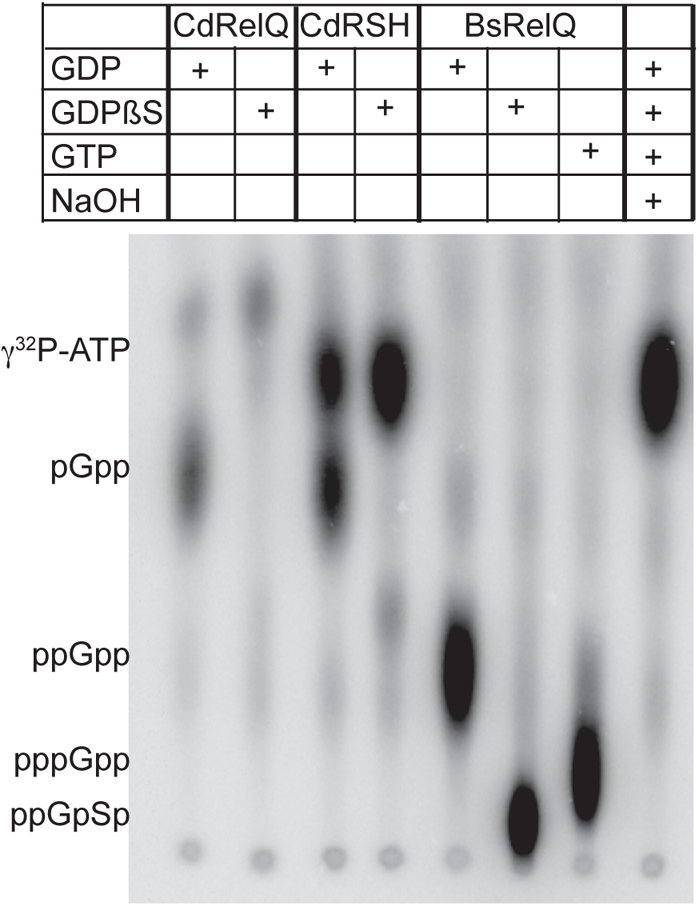
Clostridial synthetases hydrolyze the beta phosphate bond on GXP to synthesize pGpp. Phosphotransfer from γ-^32^P-ATP by 2.0 µM CdRelQ, 3.0 µM CdRSH, or 3.0 µM BsRelQ to 500 µM of GDP, non-hydrolyzable GDPβS, or GTP.

### The clostridial magic spot is pGpp

We next investigated the ^31^P chemical shifts of the (pp)pGpp synthesis reactions of both the clostridial synthetases. When ATP is mixed pair-wise with the guanosine phosphoaccepters GDP or GTP, the resonant peaks of each of the phosphorous from each nucleotide are clearly visible and are consistent with the reported values, as are the resonant peaks of AMP and GMP (Figure 6A,B, E, G).^57 58^ The chemical shifts of each of the phosphorous group are listed in Table (S2). When ATP and GDP are incubated with CdRelQ, the beta and gamma phosphate peaks of ATP at −19.16 ppm and −5.58 ppm are reduced and an AMP 5’ phosphate peak at 3.52 ppm appears, consistent with ATP hydrolysis (Figure 6A, B, C). The peaks of ATP and GDP were assigned by comparison to their peaks when analyzed individually (Figure S3). Additionally, distinct peaks at 3.52 ppm and 4.02 ppm emerge after incubation with RelQ. These are close to the 5’ phosphate peaks of AMP and GMP and may together be ascribed to the formation of AMP from ATP hydrolysis and a 5’ guanosine phosphate from hydrolysis of the beta phosphate bond of GDP (Figure 6A,C,G). We speculate that the 5’ phosphate peak of pGpp is offset from the 3.28 ppm peak of GMP by its proximity to the 3’ disphosphate moiety (Figure 6C,G). The peaks of the GDP 5’ alpha and beta phosphates, at −10.10 ppm and −6.18 ppm, are broadened, presumably by overlap with the peaks of the chemically similar pGpp 3’ alpha and beta phosphate (Figure 6B,C).

**Figure 6.**
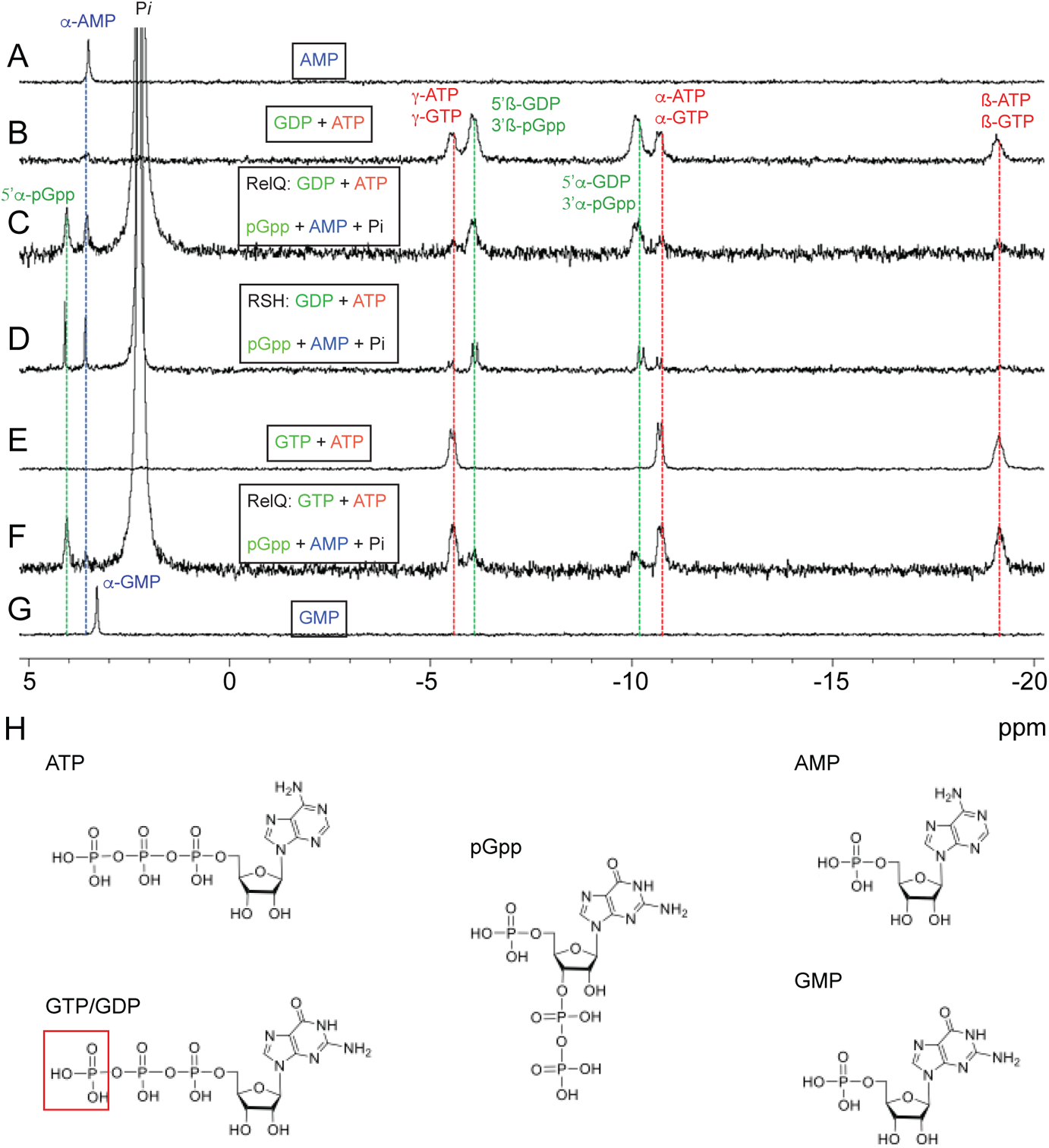
^31^P NMR evidence that clostridial magic spot is pGpp. (A) AMP standard. (B) GDP and ATP standards. (C) GDP and ATP incubated with 2.0 µM CdRelQ for 40 min. (D) GDP and ATP incubated with 3.0 µM CdRSH for 60 min. (E) GTP and ATP standards. (F) GTP and ATP incubated with 2.0 µM CdRelQ for 40 minutes. (G) GMP standard. (H) Structures of nucleotide substrates and products. GTP and GDP are differentiated by a third phosphate (outlined in red) on the 5’ hydroxyl group. The activity of the clostridial synthetases transfers a pyrophosphate from ATP to the 3’ hydroxyl of GTP or GDP, creating alpha and beta phosphate groups with chemical environments similar to those of the 5’ alpha and beta phosphates of GDP. The clostridial synthetases also cleave the 5’ beta phosphate bond of the guanosine substrate, leaving a single 5’ alpha phosphate whose chemical environment is similar to that of the 5’ alpha phosphate of GMP.

Similarly, when ATP and GDP are incubated with CdRSH, the beta and gamma phosphate peaks of ATP are almost completely abolished and the alpha and beta phosphate peaks of the GDP substrate differentiate into doublet peaks, presumably from the pGpp 3’ alpha and beta phosphates, while a pronounced peak at 4.06 ppm emerges (Figure 6B, D). We attribute the greater loss of ATP peaks and greater differentiation of the guanosine alpha and beta phosphate peaks to greater enzymatic activity by RSH, for which GDP is the preferred substrate, and subsequently less residual ATP and GDP in the reaction mixture. When ATP and GTP are mixed, the alpha and gamma phosphates of the two triphosphate substrates present as doublet peaks, while the beta phosphates present as one broad peak (Figure 6E). When ATP and GTP are incubated with CdRelQ, peaks at 4.03 ppm, −10.39 ppm, and −5.83 ppm, consistent with the 5’ phosphate and 3’ alpha and beta phosphates, respectively, emerge (Figure 6E,F).

In all of the reactions utilizing clostridial synthetases, an enormous peak is present at 2.2 ppm, identical to published values for inorganic phosphate and consistent with hydrolysis of the 5’β phosphate bond on GXP substrates by clostridial enzymes (Figure 6C,D,F). We speculate that the large inorganic phosphate peak present in the clostridial reactions shifts the 5’αP peak in the clostridial reactions away from the peak of GMP alone. ^59–60^ When the enzymes are incubated with guanosine substrates in absence of ATP, the inorganic phosphate peaks are much smaller relative to the peaks from the guanosine phosphates, suggesting that either ATP hydrolysis or formation of the 3’β phosphate bond on the alarmone product stimulate the phosphorolysis of the guanosine substrate (Figure S4). No peaks attributable to pGpp emerge, confirming that hydrolysis depends on ATP (Figure S4).

The product of the clostridial synthetases is a guanosine nucleotide containing three phosphorous atoms. Theoretically, this product could be guanosine 5’triphosphate (GTP) or guanosine 5’disphosphate-3’monophosphate (ppGp) rather than guanosine 5’monophosphate-3’disphosphate (pGpp). However, we did not observe the characteristic beta phosphate peak of GTP at −19.14 ppm in any of our NMR spectra unless exogenous GTP was provided as a substrate, demonstrating that it is not a product (Figure 6C,D,F). Additionally, the inability of the clostridial enzymes to produce any product in the absence of 5’β-phosphate bond hydrolysis eliminates ppGp as a product (Figure 5). Therefore, pGpp is the only possible product of the clostridial synthetases.

### CdRelQ utilizes diverse metal cofactors

*In vitro* analyses of RSH or SAS hydrolysis are typically carried out in the presence of micromolar magnesium. ^21, 31, 36–37, 46, 61^ RSH from *Methylobacterium exotorquens* has previously been reported to utilize cobalt more efficiently than magnesium, although its activity is severely curtailed by manganese, calcium, or nickel cofactors, indicating that the enzyme’s metal utilization is highly specific.^41^ We have previously reported that CdRSH is capable of utilizing multiple structurally diverse divalent cations as cofactors with little to no loss of efficacy compared to magnesium.^34^ Here we find that dRelQ also accepts multiple metal ion cofactors with a broad range of atomic radii, both smaller and larger than magnesium (Figure 7, Table 3). Pyrophosphotransfer to a GDP substrate is robust in almost every condition studied. GDP utilization is modestly reduced by ferrous iron, the smallest metal ion tested (Figure 7A, Table 3). CdRelQ utilization of GTP is more sensitive to metal cofactor identity. GTP utilization is decreased by iron, nickel, and copper, despite the later two being very close in size to magnesium (Figure 7A, Table 3). Even calcium, which is substantially larger than magnesium, effectively substitutes as a cofactor. In contrast, BsRelQ is much more selective for metal ion cofactors. This enzyme exhibits strong pyrophosphotransferase activity to both GDP and GTP substrates with cobalt, nickel, manganese, or magnesium cofactors, but very little activity in the presence of iron, zinc, copper, or calcium (Figure 7B). BsRelQ does appear to discriminate on the basis of ionic size, as the acceptable metal substitutions are those closest in size to the conventionally used magnesium (Figure 7B, Table 3). Nickel ions appear to disfavor the use of GTP relative to GDP by both enzymes despite the fact that nickel is very close in size to magnesium, indicating that size is not the only factor that influences metal ion cofactor specificity.

**Figure 7.**
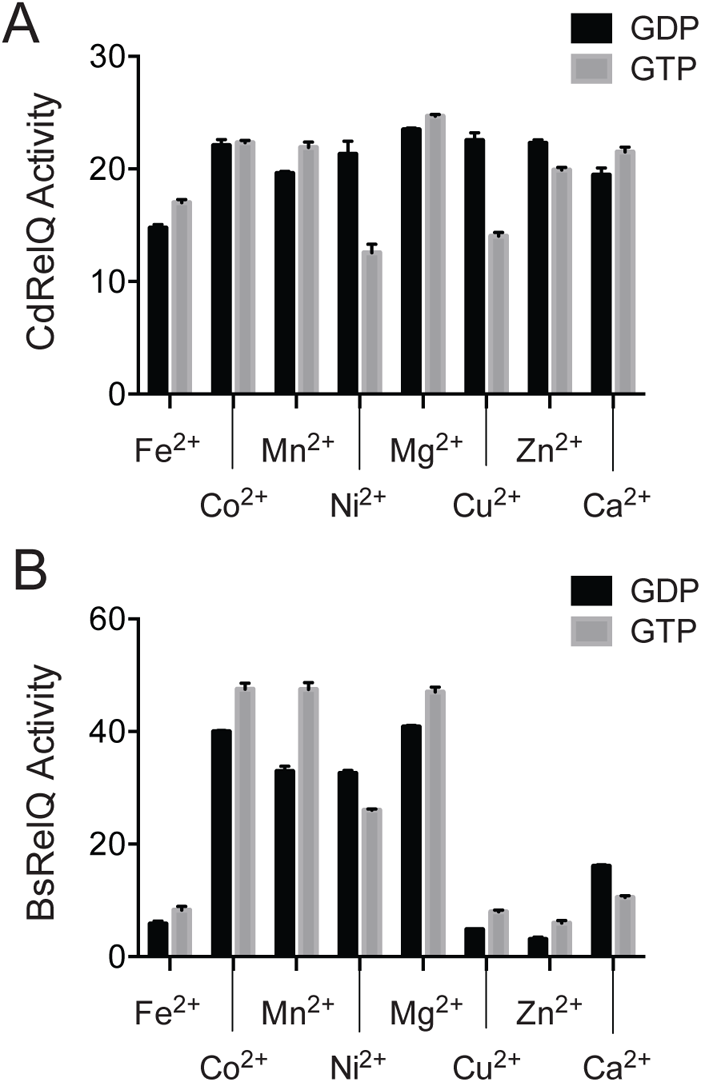
CdRelQ accommodates a wide range of metal ion cofactors. Pyrophosphotransfer after 60 minutes from γ-^32^P-ATP to 500 µM of the indicated GXP phosphoacceptor by (A) 2.0 µM CdRelQ or (B) 3.0 µM BsRelQ in the presence of 5.0 mM of the indicated metal ion cofactor. Metal ions are arranged in order of increasing ionic size.

**Table 3.** Ionic radii of divalent metal cofactors. Crystal radii of metal ions in the +2 oxidation state with octahedral coordination. Where two electron configurations are possible the low spin radius is listed.^75–76^

| Cation | Effective ionic radius (Å) |
| --- | --- |
| Fe<sup>2+</sup> | 0.75 |
| Co<sup>2+</sup> | 0.79 |
| Mn<sup>2+</sup> | 0.81 |
| Ni<sup>2+</sup> | 0.83 |
| Mg<sup>2+</sup> | 0.86 |
| Cu<sup>2+</sup> | 0.87 |
| Zn<sup>2+</sup> | 0.88 |
| Ca<sup>2+</sup> | 1.1 |

## Discussion

The outputs regulated by the stringent response are highly conserved, as accumulation of the magic spot alarmones has similar phenotypes in diverse bacteria. However, different mechanisms lead to these convergent results, and the Gram positive Firmicutes deviate in many ways from the template established by studies in γ proteobacteria such as *E. coli*. Notably, *E. coli* does not encode SAS enzymes and does not appear to use (p)ppGpp for non-SR regulation of guanosine homeostasis. Here we explore the biological role and biochemistry of the clostridial SAS RelQ. Despite the high sequence conservation between RelQ and SAS from other organisms, our results were not entirely as expected.

We have previously demonstrated that stationary phase onset and exposure to the antibiotics clindamycin and metronidazole stimulate *rsh* transcription in *C. difficile* and that genetic or chemical inhibition of RSH increases antibiotic susceptibility. ^34^ Here we confirm that antibiotic stress stimulates transcription of clostridial stringent response genes although the details of the response are strain specific; *relQ* responds to fidaxomicin but not vancomycin in R20291 and to vancomycin but not fidaxomicin in 630Δ*erm*. It is tempting to speculate that strain-specific induction of magic spot synthetase genes may contribute to observed differences in antibiotic tolerance among *C. difficile* strains, although this will need to be verified by testing in more clinical strains before a conclusion can be drawn. We have previously found that transcription of *rsh* but not *relQ* is induced by stationary phase onset. Here we report that *rsh* but not *relQ* are upregulated by oxidative and acid stress in *C. difficile* R20291. It is possible that RSH is the primary mediator of a stress-induced stringent response in this organism and that the primary role of RelQ is regulation of nucleotide metabolism as has been reported in *B. subtilis* and *E. faecalis*.^32–33^ Interestingly, our finding that medium acidification caused by stationary phase onset in R20291 is sufficient to induce *rsh* transcription in nutrient-rich fresh growth medium suggests that medium acidification due to metabolic activity may be one of the signals that regulate the expression of stationary phase genes in this organism.

Intriguingly, we report that the two clostridial magic spot synthetase enzymes share characteristics that appear to be unique to *C. difficile.* While some SAS enzymes can directly synthesize pGpp using GMP as a substrate, there is little precedent for a synthetase to produce the triphosphate signal using GDP or GTP as substrates. ^31, 40, 47–49, 62^ There are no known examples of an organism that produces pGpp exclusively. Direct synthesis of pGpp via hydrolysis of GDP or GTP 5’ beta phosphate bonds has only been reported as a minor *in vitro* activity of *E. coli* RSH, dependent on the presence of an EKDD motif in the synthetase active site.^35^ To date, every organism that synthesizes pGpp directly or indirectly also synthesizes ppGpp and pppGpp.^31, 40, 47, 62–63^ Utilization of GDP and GTP to exclusively synthesize pGpp by both a RSH and a SAS enzyme has not previously been reported. While it is still possible that RelC synthesizes the longer forms of magic spot despite its non-conserved guanosine binding motif, the dependence of CdRSH and CdRelQ on guanosine β-phosphate bond hydrolysis indicates that pGpp is a very important, and possibly the only, clostridial alarmone. It is also unusual that CdRelQ exhibits a higher affinity for GTP than GDP. The only obvious divergence between CdRelQ and characterized SAS with substrate preferences for GDP is the presence of a glycine in the first position of the GXP binding motif, which is a polar charged residue in all other SAS homologs except *C. glutamicum* RelS, which has a valine at that position and preferentially utilizes GTP (Figure 1). Previous work in RSH enzymes has posited that an acidic EXDD sequence at the beginning of the guanosine binding motif results in a preference for GDP while a basic RXKD sequence correlates to a preference for GTP. ^35^ This association does not appear to hold in SAS enzymes, several of which have a basic motif and prefer GDP (Figure 1). However, it is likely that this position does contribute to the enzyme’s ability to utilize GDP or GTP. Future mutational analysis will likely be necessary to determine what residues play a role in guanosine β-phosphate bond hydrolysis.

In addition to the unexpected triphosphate alarmone product, the characterized clostridial synthetases share another trait that does not appear to be widespread among RSH or SAS family enzymes: a remarkable ability to utilize structurally diverse divalent cation cofactors. While few other synthetases have been tested against a panel of divalent cations, cobalt, zinc, copper, nickel, iron, and calcium do not allow synthetase activity by *M. tuberculosis* Rel and manganese, nickel, and calcium reduce or abolish synthetase activity by *M. exotorquens* Rel.^39, 41^ Zinc, but not iron or nickel, stimulates enzymatic activity of crystalized RelP from *S. aureus*.^50^ There does not appear to be any precedent for the extremely broad range of metals successfully utilized as cofactors by both clostridial synthetases. ^34^ This could be a *C. difficile* adaptation to so-called ‘nutritional immunity,’ which is the ability of mammalian immune systems to chelate and sequester metal ions that are limiting factors for the growth of foreign pathogens. ^64 65^ As metal starvation can trigger the stringent response in some bacteria,^66–67^ it is possible that promiscuous utilization of divalent cations in the enzymes that initiate the clostridial stringent response could provide a survival advantage during infection.

It has been clear for some time that Gram negative *E. coli* is not a suitable model for the Gram positive stringent response. It appears that even other Firmicutes are not complete models of this pathway in *C. difficile.* Given the role of the stringent response in antibiotic tolerance and virulence, there is interest in inhibiting it as an antimicrobial strategy.^68–71^ However, the ubiquity of this signaling pathway among bacteria means that synthetase inhibitors are likely to exhibit a broad spectrum of action which could diminish the protective effect of the gut microbiota. However, the discovery that the stringent response in this organism is mediated by synthetases whose enzymatic product and metal utilization is conserved with each other but different from that of their homologs in other bacterial species raises the possibility that the clostridial enzymes share structural features that distinguish them from synthetases in other organisms.

Further characterization of the clostridial synthetase enzymes may provide a foundation for targeted inhibition of the stringent response in this organism, which could be applied as an adjuvant to antibiotic therapy to diminish *C. difficile* antibiotic survival.

## Experimental Procedures

### Cell culture

The bacterial strains and the plasmids used in this study are listed in Table S1. *E. coli* strains were grown in Luria-Bertani (LB) medium at 37°C. *C. difficile* strains were grown anaerobically in brain heart infusion medium supplemented with 5% yeast extract (BHIS). Anaerobic bacterial culture was maintained in a Coy anaerobic chamber (Coy laboratory products, Grass lake, MI) with an atmosphere of 85% N_2_, 10% CO_2_ and 5% H_2_ at 37°C. Unless otherwise stated all the experiments were conducted in *C. difficile* R20291. The pH of the *C. difficile* filtered growth medium was monitored before and after 24 hours of incubation using a benchtop pH meter (Mettler-Toledo). To avoid adding spent liquid medium to the fresh samples, cultures used for medium pH experiments were inoculated with single colonies picked from BHIS agar plates. All plastic consumables were equilibrated in the anaerobic chamber for at least 72 hours prior to use. Bacterial strains carrying plasmids were maintained using the corresponding antibiotics at the indicated concentrations: 10 µg mL^−1^ of thiamphenicol (Tm; Alfa Aesar), 50 µg mL^−1^ of kanamycin (Kan; Bio Basic).

### Promoter activity analysis

The *in vivo* promoter activity of the reporter strains was studied as previously described.^34^ Briefly, saturated starter cultures of *C. difficile* strains containing the phiLOV2.1 reporter plasmid were grown anaerobically at 37°C in BHIS-Tm10 for 12 - 16 h and inoculated 1:50 into fresh BHIS-Tm10 containing indicated treatments (1.70 µg mL^−1^ vancomycin (VWR), 0.67 µg mL^−1^ fidaxomicin (Cayman Chemical), 1.0 mM diamide (MP Biomedicals), 4.0 µM copper sulfate (Fisher Scientific)) and grown for 2 h at 37°C. To monitor temperature-induced promoter activity, log phase (OD_600_: 0.50-0.70) cells of each of the strains were divided into two halves and incubated in parallel at 37°C and 41°C for 30 mins. pH induced promoter activity was measured by inoculating exponentially growing cells 1:50 into BHIS-Tm10 containing 0.1M potassium phosphate buffer at pH 6.0, 7.0, 8.0 or 9.0. OD_600_ was recorded for each sample upon collection. To minimize discrepancies in fluorescent signal from cellular autofluorescence, cell numbers in each sample were equalized on collection. Each sample was collected at a volume that would give a cell count equivalent to 3 mL of the culture with the lowest OD_600_. After collection, cells were pelleted anaerobically in a microcentrifuge and suspended in 400 µL of anaerobic 1X phosphate buffered saline (PBS). Duplicate 200 µL samples were aliquoted into a clear-bottomed black 96-welled microplate (Thermo Fisher Scientific) and were removed from the anaerobic chamber to measure sample fluorescence intensity. Sample fluorescence using 440/30 excitation and 508/20 nm emission filters and sample OD_630_ were measured on a BrandTek plate reader. The instrumental parameters for all fluorescence measurements included a sensitivity limit of 65. Measurements were blanked against 1X PBS and were reported as E508/A630. Statistical analysis was performed using Prism (GraphPad).

### Cloning of CdRelQ

The relQ gene (CDR20291_0350) was amplified from *C. difficile* R20291 genomic DNA using primers relQ_a_F and relQ_a_R (Table S1) that added a C-terminal hexahistidine tag. The gene was subsequently ligated into the pMMBneo expression vector at the KpnI (NEB) and PstI (NEB) restriction sites, and chemically transformed into *E. coli* DH5-α first, followed by BL21. The resulting plasmid, pMMBneo::relQ was confirmed via PCR using pMMBneo plasmid-specific primers, pMMB_F & pMMB_R (Table S1). Because expression from pMMBneo did not result in a visible protein band on an SDS-PAGE gel, the relQ gene (CDR20291_0350) was amplified from *C. difficile* R20291 genomic DNA using gene specific primers (5’relQ_NdeI & 3’relQ_XhoI). The relQ amplicon was ligated into pET24a expression vector at the NdeI-XhoI restriction sites and upstream of the vector derived 6xHis-tag to generate pET24a::relQ plasmid. The plasmid was transformed into *E. coli* BL21 (New England Biolabs, NEB) and confirmed by PCR using relQ gene specific primers (relQ_a_F & relQ_a_R). (Table S1)

### In vivo overexpression of RelQ

Growth curve assays following the induction of *C. difficile relQ* in *E. coli* BL21 cells carrying pMMBneo:: relQ were performed in 96 well microtiter plates (Brand Plates) using a BioTek Synergy plate reader. Exponential phase cultures of *E. coli* were inoculated in LB medium in the presence or absence of 0.5 mM isopropyl-β-D-thiogalactopyranoside (IPTG) inducer. The plate was incubated at 37°C with shaking for a total of 16 hours while monitoring cell growth every 30 minutes.

### CdRelQ expression and Purification

Saturated starter cultures of *E.coli* BL21 cells containing pET24a-CdRelQ-His plasmid were grown overnight in LB medium with 50 µg mL^−1^ kanamycin in a shaking incubator at 250 rpm and 37°C. Cultures were diluted 1:20 into LB medium with 50 µg mL^−1^ kanamycin and grown at 37°C with shaking until the optical density at 600 nm reached 0.5-0.6. Once cells reached the desired OD_600_, they were allowed to stand at room temperature for 10 mins and induced with 0.5 mM isopropyl β-D-1-thiogalactopyranoside (IPTG) for 3 hours at 30°C for protein expression. Cells were centrifuged at 4030 rcf using a Beckman coulter JA12 rotor for 30 mins at 4°C and pellets were frozen at −20°C. Cells were lysed using lysis buffer (50 mM Tris-HCl, pH 8.0, 500 mM NaCl, 5 mM MgCl2, 10 mM imidazole, 3 mM BME, and 100 mM PMSF) via sonication. The cells were burst for 10 secs at 40% amplitude with 30 secs pause for 8 cycles followed by centrifugation at 13680×g for 15 mins at 4°C in a bench top Scilogex centrifuge. The soluble protein fraction was collected as supernatant and subjected for purification using HisPur Ni-NTA Resin affinity chromatography (Thermo Fisher Scientific). Briefly, the supernatant was mixed with equilibrating buffer (same as lysis buffer) in 1:1 ratio and subjected to 1.0 mL Ni-NTA resin column equilibrated with the equilibrating buffer. Initially the column was washed with five column volumes of the lysis buffer, and further washed and eluted using a gradient of imidazole-30 mM, 50 mM and 75 mM imidazole were used for column washing while the protein was eluted using 150 mM and 300 mM imidazole in two fractions. Purified protein was subsequently dialyzed overnight at 4°C in a buffer consisting of 30 mM Tris, pH 8.0, 300 mM NaCl, 5 mM MgCl2, 5 mM BME, and 20 % glycerol. Protein concentration was determined spectroscopically measuring A_280_ using BioDrop. Molar extinction coefficient of 36455 M^−1^cm^−1^ as determined by ExPASY^72^ for CdRelQ was used to estimate the concentration.

### Expression and purification of the control synthetase

*E. coli* BL21 cells carrying pET28 bearing the *Bacillus subtilis* SAS1/RelQ gene were generously provided by Mingxu Fang, University of California, San Diego and Carl E. Bauer, Indiana University, Bloomington. *Bacillus subtilis* RelQ (BsRelQ) was expressed and purified using a protocol developed by Mingxu Fang as previously published.^34^ Briefly, *E. coli* BL21 cells bearing pET28a::BsRelQ were grown in LB media containing 50 µg mL^−1^ kanamycin at 37°C with shaking until the OD_600_ reached 0.4. Protein expression was induced overnight at 16°C by the addition of 200 µM IPTG. Cells pellet was collected by centrifugation at 4°C and resuspended in StrepTactin buffer containing 50 mM Tris pH 8.9, 1 M NaCl and 20% glycerol along with 2 mM PMSF. The cells were sonicated with 10 secs burst and 30 secs pause cycle at 40% amplitude for 8 cycles followed by centrifugation to collect the clarified lysate. The protein was purified by using StrepTactin resin column (IBA Life sciences) following the manufacturer’s instructions.

### In vitro measurement of synthetase activity

(pp)pGpp synthesis was carried in a reaction volume of 10.0 µL as described previously^34, 73^ with a fixed concentration of CdRelQ or BsRelQ. The reaction mixture consists of a buffer containing 10 mM Tris-HCl (pH 7.5), 5 mM ammonium acetate, 2 mM KCl, 0.2 mM DTT, 0.15 mM ATP, 5mM MgCl_2_, 1.0 µCi of γ-^32^P-ATP (PerkinElmer), and indicated concentrations of AMP (Acros organics)/ADP/ATP (Bio Basic)/GMP (BioPlus Chemicals)/GDP/GTP (BioBasic) phosphoacceptor. Where indicated, the pH of the reaction was adjusted with HCl or NaOH. Where indicated, MgCl_2_ was substituted with 5 mM metal salts in the +2 oxidation state (MnCl_2_, CoCl_2_, CuCl_2_, ZnCl_2_, NiBr_2_, CaCl_2_, and FeSO_4_). The reactions occurred at 37°C and were stopped by spotting 2.0 µL of each sample onto PEI-cellulose TLC plates, allowing the spots to dry. Unless otherwise indicated, reactions were run for 60 minutes before quenching. The plates were subsequently developed in 1.5 M KH_2_PO_4_ (pH 3.64 ± 0.03) and autoradiographed using a Storm 860 Phosphorimager (GE Healthcare Life Sciences). pGpp signal was quantitated using ImageJ software.^74^ RelQ activity was expressed as the percentage of γ-^32^P-ATP substrate converted into product at each timepoint.^73^ Michaelis-Menten assays were performed by running the reactions for 5 minutes at the indicated concentration of GDP or GTP. Data shown are the mean and standard deviation of two experiments. The line of best fit for the Michaelis-Menten equation was calculated using non-linear regression with least squares fit and the equation y = (V_max_*X)/(K_M_ + X) where X represents the GXP substrate concentration and y represents the initial reaction velocity and graphed with Prism (GraphPad).

### Hydrolase contaminant check

BsRelQ synthesis reactions containing either GDP or GTP as substrates were conducted for 60 minutes at 37°C as described above. CdRelQ was added to each reaction at a final concentration of 2.0 µM and incubated for an additional 60 minutes. At 0, 30, and 60 minutes after CdRelQ addition, 2.0 µL of each reaction were spotted into the PEI-TLC plates for development as described above.

### GXP phosphatase activity

Guanosine nucleotide hydrolysis was assessed through synthetase assays performed as above using guanosine 5’-β-thio-diphosphate trilithuim salt (GDPβS) as the only nucleotide phosphoacceptor (Millipore Sigma). The reactions were caried out for 60 mins at 37°C and the spots were autoradiographed as reported above.

### 31P NMR spectroscopy

The structural properties of the newly synthesized product were also evaluated by phosphorous NMR. Synthesis reactions were carried out with clostridial synthetases as mentioned above for 40 mins at 37°C and the reaction mixtures were subjected to acquire phosphorous NMR spectra. In addition, phosphorous NMR spectra of the standard nucleotides (ATP, GTP, GDP, GMP, AMP) solutions in the reaction buffer were also acquired for signal comparison and assignments. D_2_O at 15% was added into each of the samples to enable locking of the magnetic field in the NMR spectrometer. NMR spectra were indirectly referenced to 85% external phosphoric acid. ^31^P NMR recordings with proton decoupling were performed using 400 MHz Bruker Avance III HD NMR spectrometer at 162 MHz (^31^P) equipped with 5 mm PABBO BB/19F-1H/D Z-GRD Z108618/0798 probe. A total of 14000 scans for each sample at 303K were performed to acquire NMR spectra. All the measurements were carried out using pulse sequences supplied by the spectrometer manufacturer (Bruker-TopSpin 3.2).

### Isothermal calorimetry (ITC)

ITC200 microcalorimeter (Malvern) was employed for the analysis of the binding affinity of CdRelQ towards GXP as described previously.^34^ Briefly, a total volume of 300 µL reaction buffer containing 0.006 mM of the protein was added into the cell component and 40 µL of 0.06 mM GXP was filled into the syringe from which a total of 38 injections were made for the analysis at 37°C. Both the protein and ligands were prepared in the same buffer containing 10 mM Tris-Cl, 5 mM AmAce, 5 mM MgCl_2_, 0.2 mM DTT, 0.12 mM ATP and 2 mM KCl. The data obtained were processed in Origin software using a single-binding-site model. The heats of dilution acquired through the titration of ligand into the reference solution was subtracted from the binding curves during peak integration and the calculation of thermodynamic parameters. Two independent samples were measured.

## Acknowledgements

We thank Dr. Carl Bauer of Indiana University, Bloomington for the BsRelQ expression vector and Dr. Mingxu Fang of University of California, San Diego for very helpful troubleshooting discussions during enzyme purification. We thank Dr. Alvin Holder of Old Dominion University for use of the ITC instrument and helpful discussion of metal cofactors, and Dr. Michael Celestine of Old Dominion University for assistance with ITC. We thank Dr. Steven Pascal of Old Dominion University for very helpful discussion of NMR data.

The data that support the findings of this study are available from the corresponding author upon reasonable request. The TLC image in Figure 4B was previously reported in Appendix D of Astha Pokhrel’s doctoral dissertation ‘Evaluating the Role of the Stringent Response Mechanism in *Clostridioides difficile* Survival and Pathogenesis.’

This work was funded by the National Institutes of Health 1K22AI118929-01 and by start up funds from Old Dominion University. Estevan J. Coronado and Marrett M. Gilfus were supported by NSF REU CHE-1659476. The authors declare no financial conflict of interest.

---

**Figure S1.**
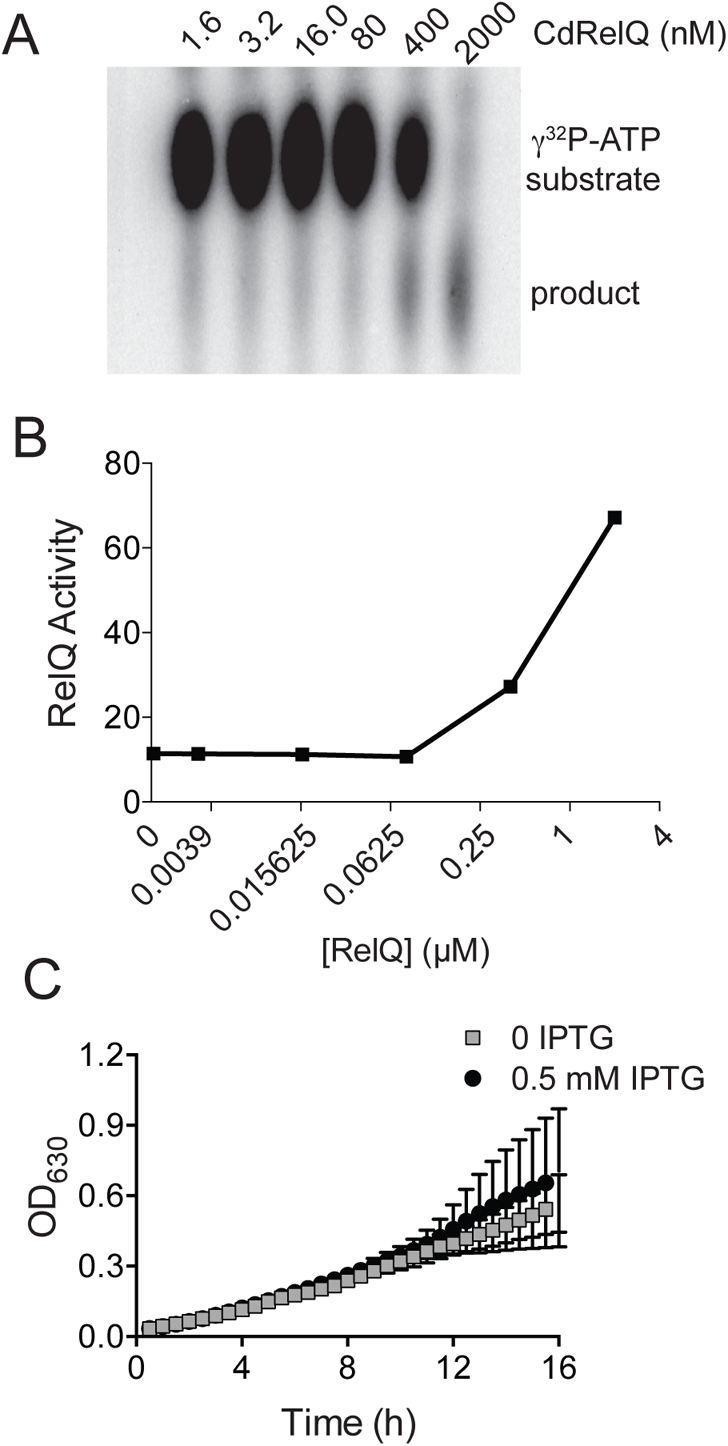
CdRelQ activity *in vitro* is concentration-dependent. (A) A representative activity assay shows transfer of pyrophosphate from γ-^32^P-ATP to a GDP acceptor requires a high concentration of RelQ. (B) Quantification of pyrophosphotransfer activity shown in panel A. (C) Growth of *E. coli* BL21 carrying pMMBneo::CdRelQ-His6 in the absence of presence of 0.5 mM IPTG inducer. Shown are the means and standard deviations of two biologically independent samples measured in triplicate, with one contaminated sample excluded from analysis.

**Figure S2.**
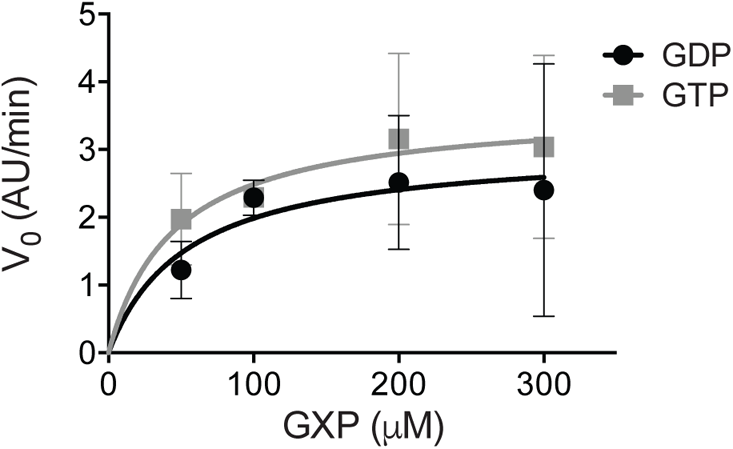
CdRelQ substrate affinity. Michaelis-Menton plot showing the initial reaction velocity of CdRelQ pyrophosphotransfer to the indicated concentrations of GTP or GDP.

**Figure S3.**
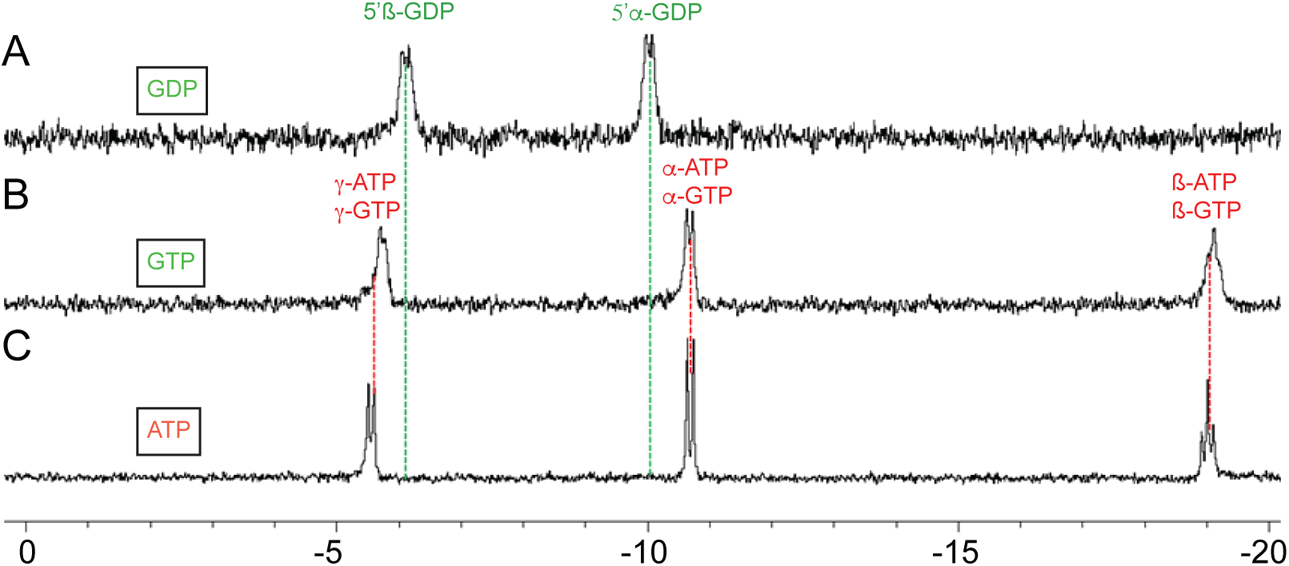
NMR spectra of nucleotide standards. 31P NMR spectra of (A) GDP, (B) GTP, and (C) ATP.

**Figure S4.**
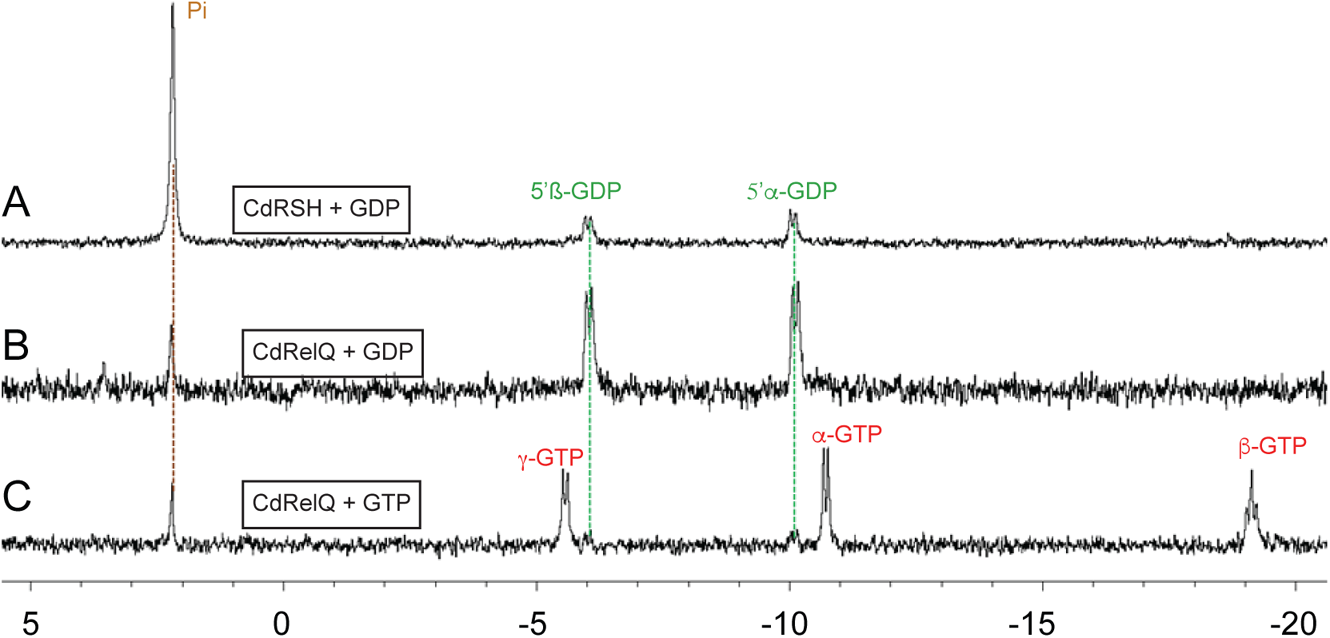
^31^P NMR of clostridial synthetase reactions without ATP. (A) GDP incubated with 3.0 µM CdRSH for 40 min. (B) GDP incubated with 2.0 µM CdRelQ for 40 min. (C) GTP incubated with 2.0 µM CdRelQ for 40 min.

**Table S1:**
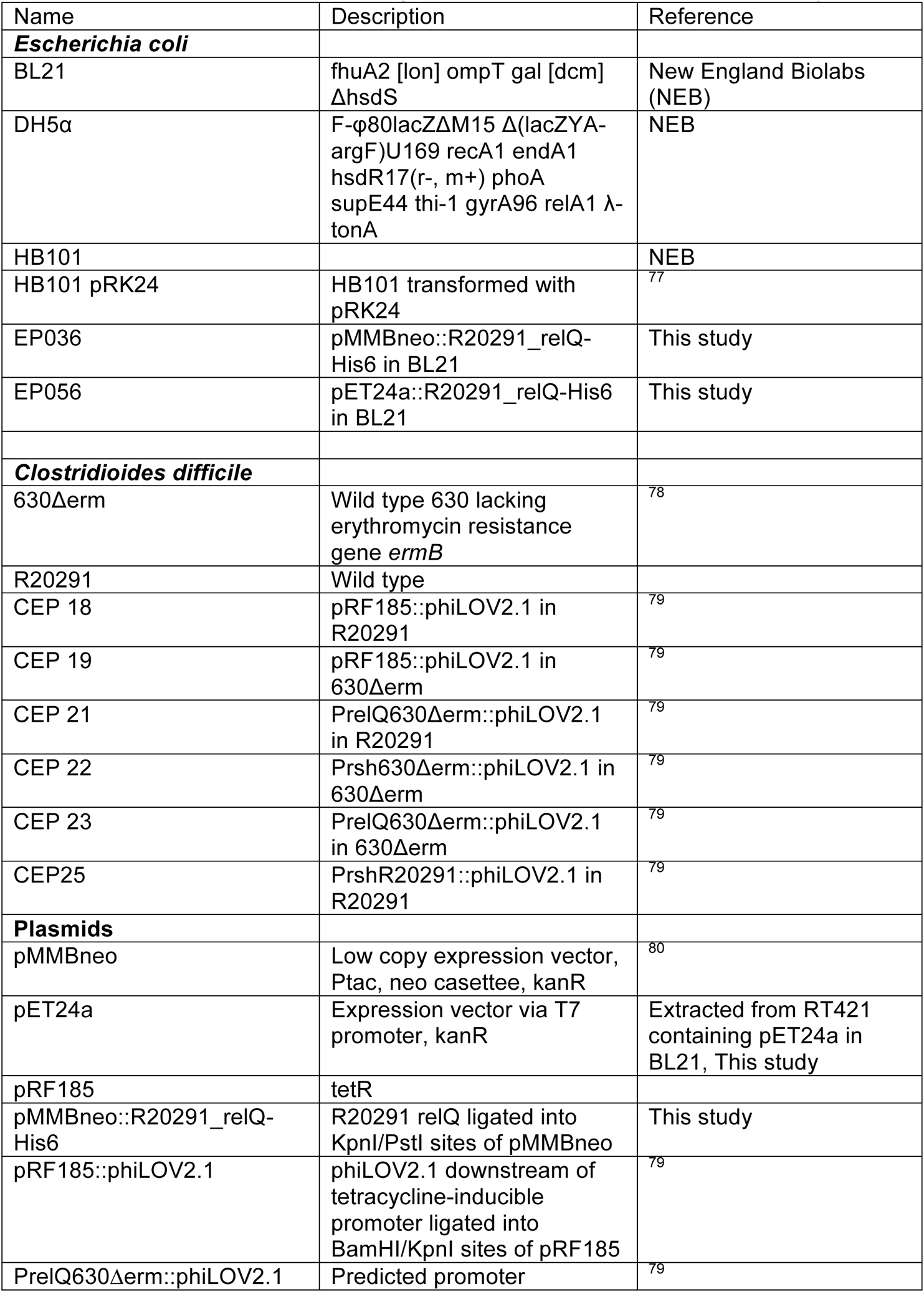
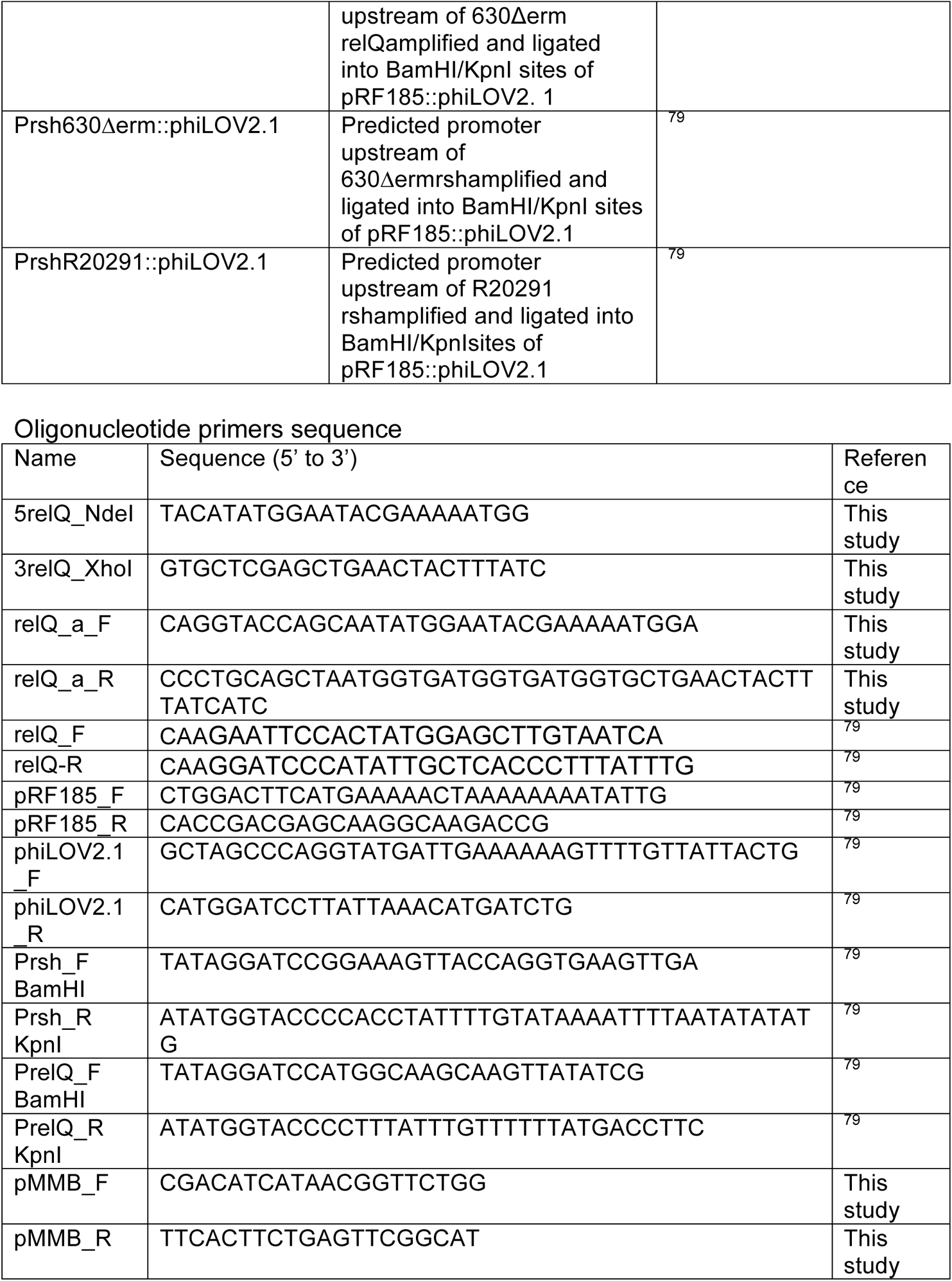
Strains, plasmids and oligonucleotide primers used in this study Oligonucleotide primers sequence

